# Multidimensional molecular differences between artificial and wild *Artemisia rupestris* L.

**DOI:** 10.1101/2021.02.26.433081

**Authors:** Zhi Zhou, Bin Xie, Bingshu He, Chen Zhang, Lulu Chen, Zhonghua Wang, Yanhua Chen, Zeper Abliz

## Abstract

Different ecological environments affect the active ingredients and molecular content of medicinal plants. *Artemisia rupestris* L. is a kind of traditional medicinal plant, and the shortages of the wild resource have led to increased use of artificial varieties. However, there have few investigations referring to molecular differences between them in a systematic manner. In the present study, artificial and wild *Artemisia rupestris* L. plants were collected in the Altay–Fuyun region, Xinjian, China. Untargeted metabolomics method based on liquid chromatography-mass spectrometry (LC-MS) technology was applied to profile flower, stem, and leaf samples, respectively, and levels of a panel of representative known metabolites in this plant were simultaneously analyzed. The genetic basis of these samples was explored using a *de novo* transcriptomics approach to investigate differentially expressed genes (DEGs) and their pathway annotations. Results indicated metabolic differences between the two varieties mainly reflected in flavonoids and chlorogenic acid/caffeic acid derivatives. 34 chemical markers (CMs) belonging to these two structural categories were discovered after validation using another batch of samples, including 19 potentially new compounds. After correlation analysis, total of six DEGs in different organs relating to 24 CMs were confirmed using quantitative real-time PCR (qPCR). These findings provided novel insight into the molecular landscape of this medicinal plant through metabolomics-transcriptomics integration strategy, and reference information of its quality control and species identification.

**ONE SENTENCE SUMMARY:** A metabolomics-transcriptomics research on Artemisia rupestris L. to discover metabolite differences and the genetic basis between artificial and wild varieties in systematic and novel manner.

## INTRODUCTION

*Artemisia rupestris* L. (compositae, sagebrush) is a plant used in traditional Chinese Uyghur medicine and is mainly found in the Xinjiang region of China, central Asia, and Europe (Gu et al., 2012). It has anti-inflammatory, antibacterial, antivirus, anti-allergy, antitumor, and liver protective properties (Liu et al., 1986; Xiao et al., 2008; Guo et al., 2009; Fang et al., 2011). It is commonly used in Uyghur medicine as whole herb, and it is also the key ingredient in Compound Yizhihao granule that used to treat colds in China (Gu et al., 2012). Because of its unique ecological characteristics, *Artemisia rupestris* L. mainly depends on wild resources. However, increased demand and overdevelopment in recent years has depleted this resource and artificial cultivation has gained increasing attention.

To date, more than 100 components have been reported in *Artemisia rupestris* L. The active ingredients mainly include flavonoids, sesquiterpenoids, organic acids, and alkaloids (Liu et al., 1985; Song et al., 2006; Ji et al., 2007; Su et al., 2008; Su et al., 2010; Gu et al., 2012; He et al., 2012). Rupestonic acid, a type of guaiacanesequiterpenoid, is a characteristic metabolite present in *Artemisia rupestris* L. that clearly shows anti-influenza virus activity. Various rupestonic acid derivatives have been synthesized to screen for agents with improved anti-influenza activity and lower toxicity (Zhou et al., 2012; He et al., 2014; Zhao et al., 2017). Overall, previous studies into *Artemisia rupestris* L. mainly focused on partial high-abundance components or specific categories of secondary metabolites. While, as a complex organism, the whole metabolome, including primary and secondary metabolites, may be involved in its pharmaceutical effect. There is still a lack of comprehensive and global understanding about the material basis of this medicinal plant. Furthermore, plant samples are usually treated after drying naturally, which may alter the content of its constituents and lead to inaccurate assessment.

Metabolomics is the systematic study of small molecular metabolites in biological samples under particular physiological or pathological conditions (Nicholson et al., 1999; Fiehn et al., 2000). It has been used in plants to investigate conditions such as abiotic stress responses (Tiago et al., 2016), diversity (Kusano et al., 2015), evolution (Xu et al., 2019), and growth (Wei et al., 2017). Since plants, including medicinal plants, cannot escape environmental conditions that adversely affect their growth and development, thus understanding fluctuations in their metabolome resulting from different ecological factors is important to their pharmaceutical effect and quality control. At present, few metabolomics studies have examined *Artemisia rupestris* L. We previously established a metabolomics analytical method for this plant (Chen et al., 2018) and applied to investigate differences between samples from Altay–Fuyun and Hami regions (Xie et al., 2020), which provided methodological basis for subsequent studies.

Multi-omics integration approaches have already been applied due to the rapid progress in high-throughput data generation (Jamil et al., 2020). Compared with studies using metabolomics methods independently, other “omics” data could provide a deeper interpretation and enhance our understanding of the variation in metabolites. In plants, integrating multiple omics methods remains challenging, largely due to poorly annotated genomes, multiple organelles, and diverse secondary metabolites (Jamil et al., 2020). Transcriptomics studies focus on the expression levels of the transcriptionally active elements within genomes (de Wit et al., 2012), and still a good option even in the absence of a reference genome (Haas et al., 2013).

In the present study, a combined untargeted and targeted metabolomics strategy was used to analyze flower, stem, and leaf samples from *Artemisia rupestris* L. to examine the material differences between artificial and wild varieties, and identify reliable chemical markers (CMs). Simultaneously, a *de novo* transcriptomics method was adopted to investigate the genetic basis of differences between the varieties using matched samples. Pathway and correlation analyses were used to screen candidates differentially expressed genes (DEGs), which were then confirmed by quantitative real-time PCR (qRT-PCR). The present study was designed to examine the molecular differences between the artificial and wild varieties of *Artemisia rupestris* L. on multiple levels, and provide a strategy reference for phytomedicine studies.

## RESULTS

### Establishment of a metabolomics–*de novo* transcriptomics integration strategy to compare *Artemisia rupestris* L. varieties

A strategy integrating metabolomics and transcriptomics methods was first established to systematically identify the molecular differences between artificial and wild *Artemisia rupestris* L. (Fig.1). Flower, stem, and leaf samples were analyzed separately based on our previous findings that revealed significant differences in the metabolome of the different organs in this plant (Chen et al., 2018). An untargeted metabolomics method was applied to profile global situations of two groups; simultaneously, a panel of known representative metabolites was targeted analyzed in case important information was missing. Differential metabolites consisting of unknown and known compounds were validated using another batch of samples to obtain reliable CMs. *De novo* transcriptomics analysis was conducted due to an absence of a reference genome. Differences between groups could be observed on a genetic level, and key DEGs involved in related metabolic pathways of CMs could be identified based on their structure classification. Correlation analysis was used to screen for candidate DEGs for further validation and qRT-PCR was used to confirm and interpret the genetic basis of CMs.

### Metabolic profiling comparison between artificial and wild *Artemisia rupestris* L.

To ensure reliable results, the quality of data was examined using randomly arranged real samples, testing stability monitoring of deviation of quality control (QC) samples, and separation performance stability assessment using retention time deviation of all samples. Subsequent analyses were performed under conditions that the data quality met the requirements (Zhou et al., 2017). Typical total ion chromatograms (TICs) revealed that although the metabolic profiles were similar between the artificial (A) and wild (W) groups and groups regardless of the organ, several significant differences were detected (Fig.2A). Initial analyses revealed retention time regions between 12 and 15 min, and 19 and 22 min for flower samples, some metabolites responded much higher or were only present in one variety. Furthermore, samples from two groups were completely separated in the score plots of the unsupervised principal component analysis (PCA) models.

**Figure 1.**
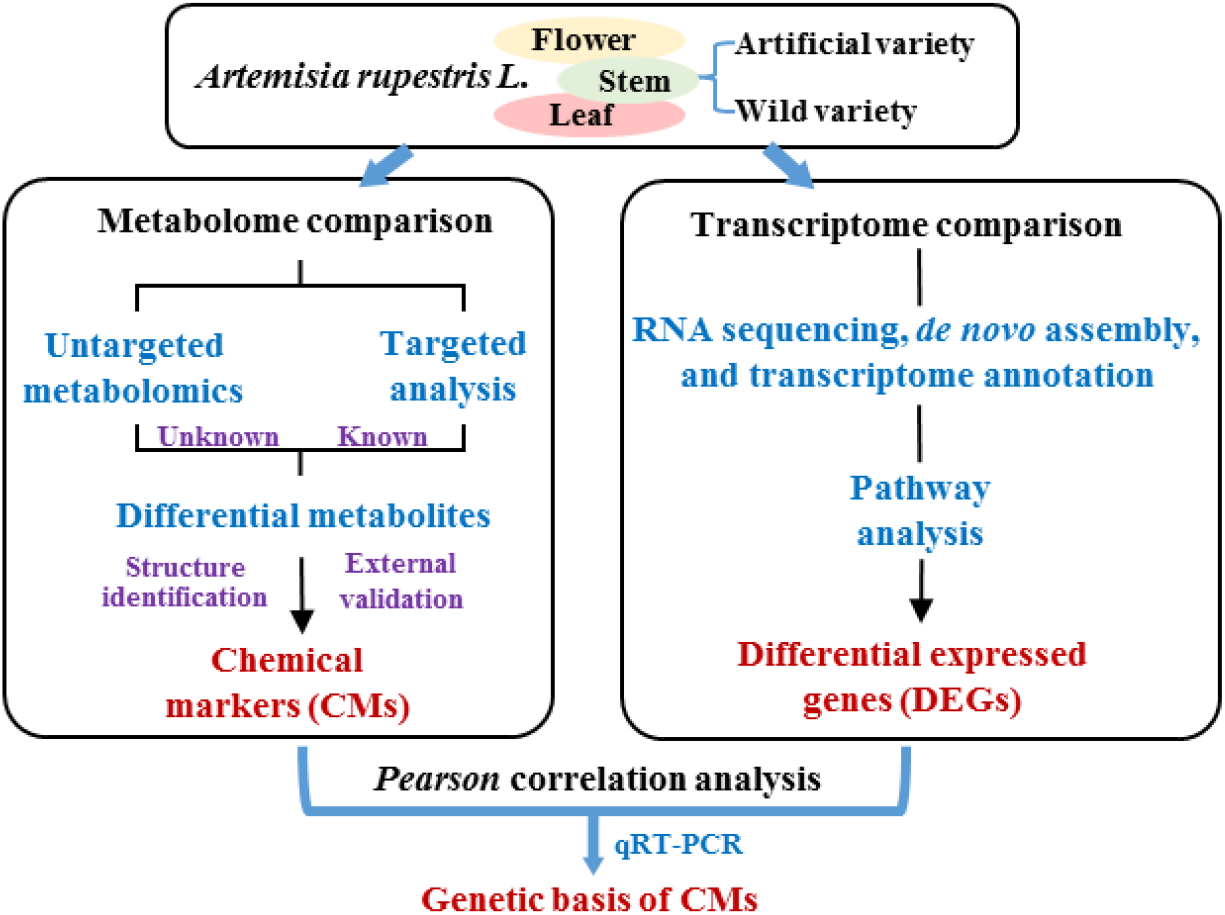
Scheme of the metabolomics–*de novo* transcriptomics integration strategy for molecular differences mapping in artificial and wild *Artemisia rupestris* L. and chemical marker discovery

Supervised orthogonal partial least square-discriminate analysis (OPLS-DA) models were established after validation of effectiveness, predictive capacity validation, and permutation in order to screen for the differential metabolites contributing to group separation. Filtration of differential variables was subsequently performed, and only those with a variable importance parameter score (VIP) >1 were considered. Adduct and isotope ions, as well as variables with crossover between groups, were also deleted. The number of differential variables and those with a fold change (FC) >10 in the three organs are shown in Fig.2B. The total numbers of differential variables in the three organs were close; however, the number of variables with FC >10 in the leaf sample was the highest.

**Figure 2.**
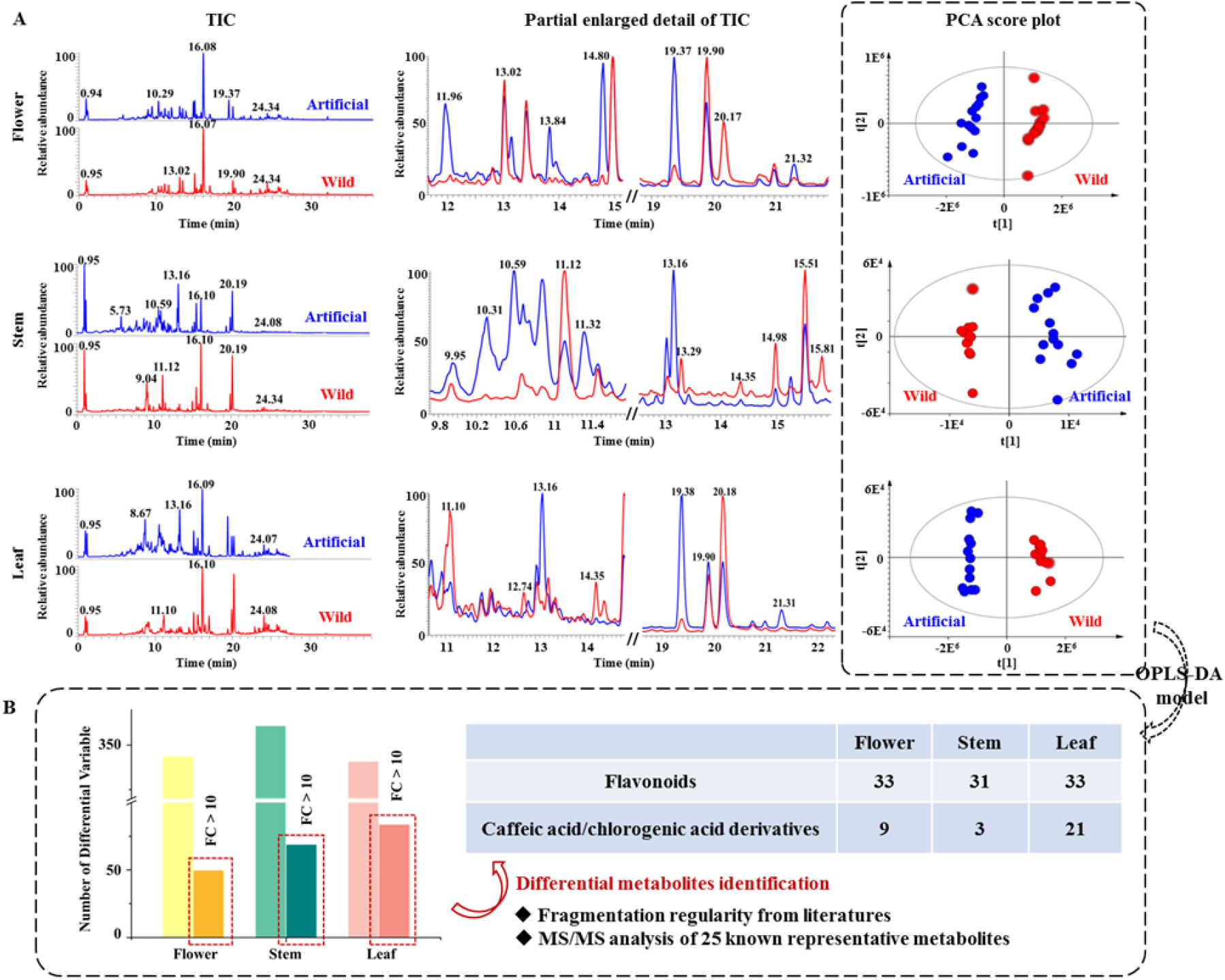
Comparison of the metabolic profiling between artificial and wild *Artemisia rupestris* L. based on ultrahigh performance liquid chromatography–electrospray ionization–mass spectrometry (UHPLC-ESI-MS) data. (**A**) TICs and partial enlarged details comparison between the two groups of flower, stem, and leaf samples, and corresponding PCA score plots. (**B**) The number of differential variables and structural categories of known differential metabolites between the two groups in the three organs.

Total of 25 known representative metabolites were targeted analyzed, and 18, 16, and 11 were detected in the flower, stem, and leaf, respectively (Supplemental Table S1). Combined with univariate statistical analysis and the variable plot of the original data, 11, 6, and 9 known metabolites showed significant differences between groups in the three sample types. The results were similar among the different organs, and mostly had FC <5 (Fig. 3). In particular, linarin (M24) and rutinum (M25) showed a higher abundance in the W group. Analysis of individual organs revealed isoquercitin (M21) and hyperoside (M22) in the A group flower, caffeic acid (M2) and apigenin (M6) in the W group flower, acacetin (M7) in the W group stem, isohamnetin (M12) in the A group leaf, and chlorogenic acid (M14) in the W group leaf were more than five times higher than in their counterparts. Rupestonic acid showed no significant difference between the two groups, but with a slightly higher content in the A group.

**Figure 3.**
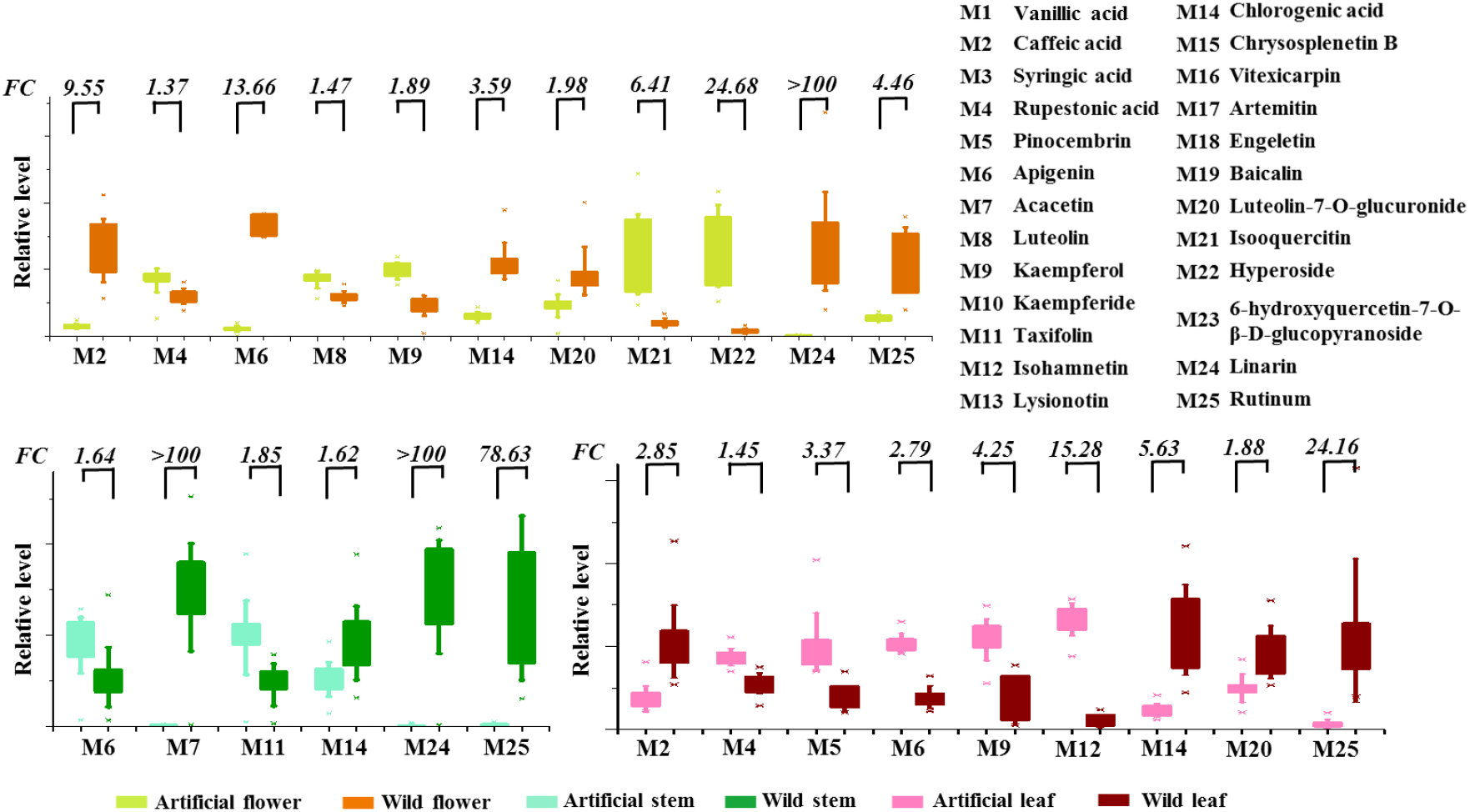
Evaluation of representative known metabolites between artificial and wild *Artemisia rupestris* L. in flower, stem, and leaf.

### Structural identification of differential metabolites

Structural classification of differential variables with FC >10 and qualified MS/MS data were conducted combining the analysis of the 25 known metabolites and retrieval of the cleavage law literature of the same categories.

Flavonoids have obvious mass fragmentation regulars. For example, loss of C_3_O_2_ (−67.9893) would occur in flavones or isoflavones with 5,7-dihydroxyl substituted A-ring and hydroxyl substituted B-ring to product [M−H−C_3_O_2_]^−^ (Kang et al., 2007). Additionally, isoflavones may eliminate CO_2_ (−43.9893), CO (−27.9944), or CHO· (−29.0022) at the C-ring via a seven-membered structure, while the C-ring in flavones would lose C_2_H_2_O (−42.0103) to generate [M−H−C_2_H_2_O]. In the case of *O*-methylated flavonoids, loss of 15 amu (−CH_3_) is common. Furthermore, the ^1,3^A^−^ ion (the fragment ion originating from 1/3 bond cleavage in the C-ring, and containing an intact A-ring) derived from the retro-Diels-Alder reaction is observed for all flavonoid subclasses (Cuyckens et al. 2004; de Rijke et al. 2006). In terms of glycosylation compounds, aglycone ions and aglycone radical ions are likely to be present in the MS/MS spectra (March et al, 2004; Vukics et al, 2010).

The MS behavior of chlorogenic acid derivatives was also analyzed. The main product ions were at *m/z* 191.0538 [quinic acid−H] and *m/z* 179.0538 [caffeic acid−H], with an occasional presence of *m/z* 146.0901 [cinnamic acid−H]or *m/z* 193.0501 [ferulic acid−H]. The rule that 191 appears as the base peak was validated as a common feature of chlorogenic acid with acyl groups connecting to 3-OH or 5-OH on quinic acid (Clifford et al. 2005; Gouveia et al. 2010). The ion at *m/z* 173.0384 [quinic acid−H_2_O−H] is likely to be the base peak in the MS/MS spectra of compounds with acyl groups linked to the 4-OH position of quinic acid.

Analysis of the MS/MS spectrum of rupestonic acid revealed that the [M−H]−ion at *m/z* 247.1334 generated high-abundance *m/z* 203.1436 by losing CO_2_ (−43.9898), which could be a diagnostic ion corresponding to the acid skeleton. Furthermore, ions at *m/z* 163.1123 and *m/z* 135.0839 presented at higher collision energy were obtained from further cracking of *m/z* 203 (Gu et al. 2012). Thus, metabolites with product ions at *m/z* 247, 203, 163, and 135 in their MS/MS spectra were recognized as rupestonic acid derivatives.

Finally, 97 flavonoids (including 33, 31, and 33 in flower, stem, and leaf samples, respectively), 35 chlorogenic acid/caffeic acid derivatives (including 9, 3, and 21 in flower, stem, and leaf samples, respectively) were classified (Fig. 1B).Rupestonic acid derivatives were also identified and all had FC values <10, with the majority <5.

### RNA sequencing, *de novo* assembly, and transcriptome annotation in *Artemisia rupestris* L.

We sequenced RNA libraries derived from flowers, stem, and leaves. Illumina sequencing from RNA-sequencing libraries yielded 39.7 million to 45.3 million reads, with 39.8%–44.2% GC content obtained after eliminating the adaptor sequences. Fast QC analysis showed that 85.78%– 91.79% of the total sequences were of quality >Q30. Since the genome sequence of *Artemisia rupestris* L. plant is not available, the reads were further assembled *de novo* into a total of 439,612 unigenes with N50 of 1428 bp (Supplementary Table S2).

Functional annotation of the 439,612 unigenes was performed by searching public databases. Overall, 185,724 (42.3%) unigenes were annotated according to their similarities with known genes/proteins. The assembled transcripts were annotated using BLASTx against the Nr database. The match result showed that the top hits for 46.81% of unigenes were from *Cynaracardunculus* var. *scolymus* (46.81%), followed by *Vitisvinifera* (2.49%), and *Hordeumvulgare* subsp. *vulgare* (1.97%) (Fig. 4A). Gene ontology (GO) analysis (Fig.4B) identified biological process, metabolic process, and cellular process were the dominating terms followed by single-organism process. Among the cellular component, 35.56%, 35.16%, and 31.37% of the annotated genes were classified into the GO terms cell, cell organ, and membrane. In the molecular function group, catalytic activity and binding were the principal GO terms.

**Figure 4.**
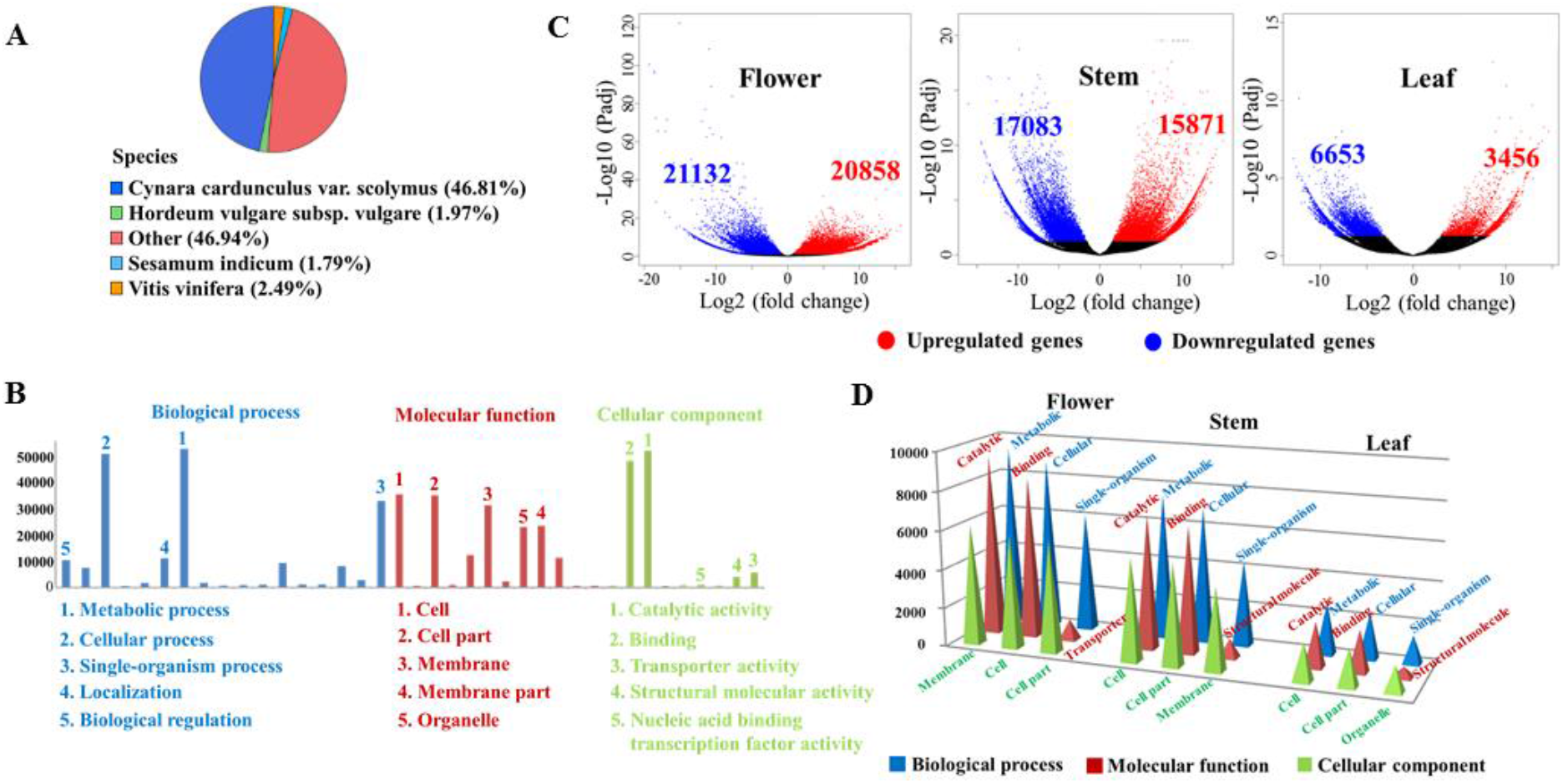
Overview of *de novo* transcriptomics analysis of *Artemisia rupestris* L. (A) Top blast species distribution for BLASTx matches. (B) GO assignment of unigenes. (C) Volcano plot of unigenes expression level in flower, stem, and leaf samples. (D) Top GO function classification statistics of differentially expressed unigenes in flower, stem, and leaf samples.

### Identification of DEGs

The expression profiles of separated organs from two varieties *Artemisia rupestris* L. were analyzed (Fig. 4C). In flower samples, 20,858 unigenes were upregulated in the W group and 21,132 were downregulated. There was a total of 32,954 (15,871 upregulated and 17,083 downregulated in the W group) and 10,109 (3,456 upregulated and 6,653 downregulated in the W group) DEGs in the stem and leaf, respectively. All DEGs were subjected to GO analysis (Fig. 4D). In the biological process category, the top three processes of DEG assignment for the three organs were the same (metabolic, cellular, and single-organism processes). However, the number of DEGs in each process differed, with 6104–9360 in flower samples, 4374–7460 in stem samples, and 1452–2670 in leaf samples. “Catalytic activity” and “binding” were the first two in molecular function terms, followed by transporter activity for flower samples and structural molecules for stem and leaf samples.

DEGs were subjected to KEGG pathway enrichment analysis. In flower samples, 21,569 unigenes had hits for 135 KEGG pathways. According to enrichment index Q value (false discovery rates corrected), “starch and sucrose metabolism” containing 743 DEGs was the top item, followed by “phenylpropanoid biosynthesis,” “other glycan degradation,” “plant–pathogen interaction,” and “stlbenoid, diarylheptanoid and gingerol biosynthesis.” In stem samples, 17,236 unigenes were assigned to 135 KEGG pathways, and the top five enriched pathways were “ribosome,” “oxidative phosphorylation,” “biosynthesis of unsaturated fatty acids,”“alpha-linolenic acid metabolism,” “circadian rhythm-plant.” In contrast, in leaf samples, 4,452 unigenes were recognized as items in 134 KEGG pathways, in which “ribosome,” “oxidative phosphorylation,” “biosynthesis of unsaturated fatty acids,” “alpha-linolenic acid metabolism,” and “fatty acid metabolism” were at the top of the list.

### External validation for CM discovery

Differential metabolites originating from the multivariate statistical and targeted analysis were validated using another batch of samples to confirm their reliability. In this assay, a targeted analysis method based on parallel reaction monitoring (PRM) was established, which is described as high-resolution multiple reaction monitoring (Ronsein et al., 2015). LC–MS parameters were the same as those used in the untargeted analysis, with the collision energy values optimized according to the character of each metabolite. Among the differential metabolites, total of 34 were confirmed as CMs. These CMs could be completely and correctly distinguish in the A and W groups (Fig. 5), indicating their high effectiveness and reliability. The majority of the metabolites had a much higher content in the W group, and the FC values ranged from 10 to several thousands. The 34 CMs included 11 chlorogenic acid/caffeic acid derivatives (CCDs), 23 flavonoids, and their putative identification is shown in Table 1 (detailed MS/MS data can be found in Supplementary Table S3). SciFinder retrieval identified 19 of the CMs as newly reported compounds due to their absence in the database.

**Table 1.**
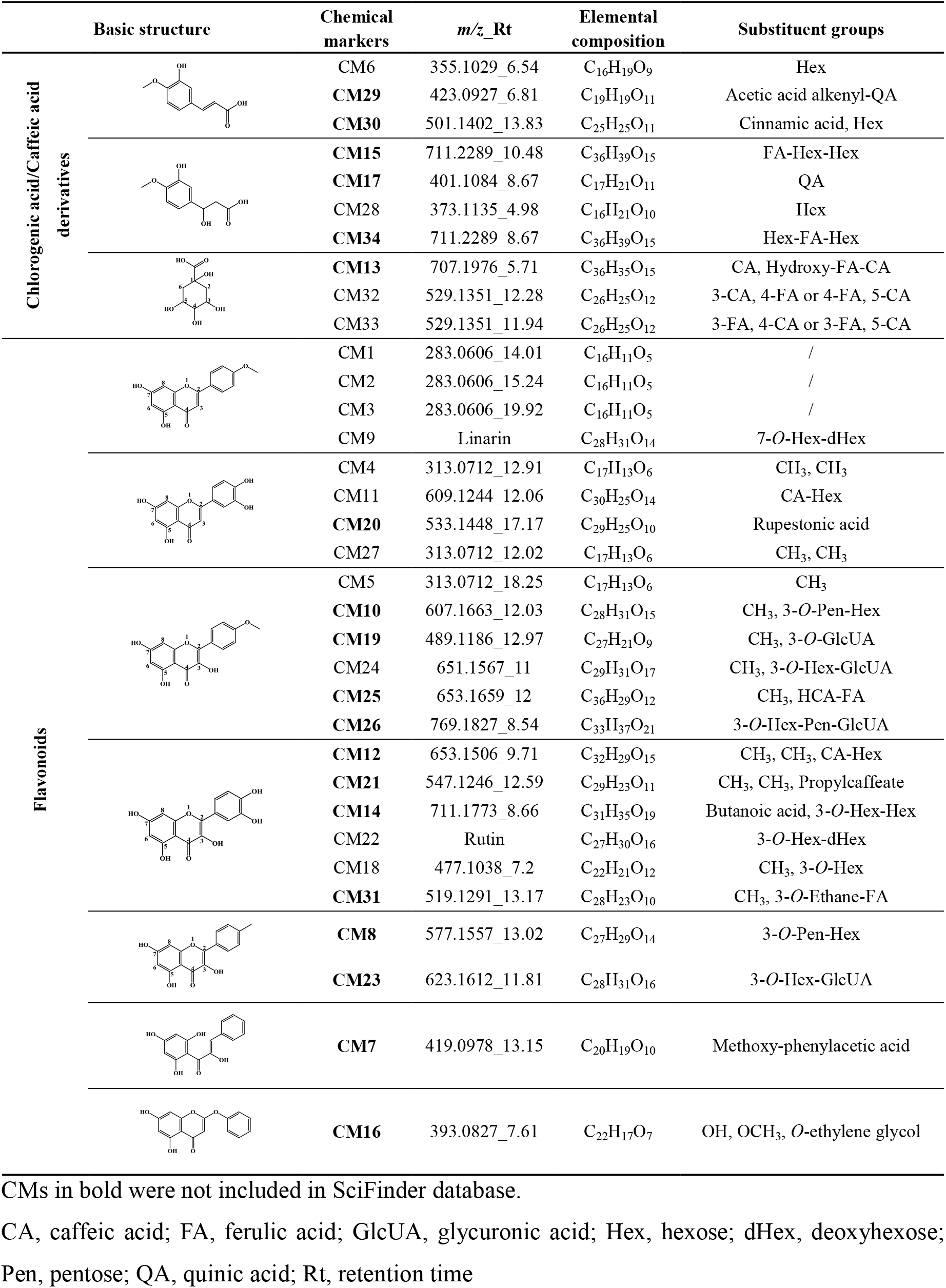
Structural identification of 34 CMs in artificial and wild *Artemisia rupestris* L. CMs in bold were not included in SciFinder database.

|  | Basic structure | Chemical markers | *m/z*_Rt | Elemental composition | Substituent groups |
| --- | --- | --- | --- | --- | --- |
| Chlorogenic acid/Caffeic acid derivatives |  | CM6 | 355.1029_6.54 | C<sub>16</sub>H<sub>19</sub>O<sub>9</sub> | Hex |
|  |  | CM29 | 423.0927_6.81 | C<sub>19</sub>H<sub>19</sub>O<sub>11</sub> | Acetic acid alkenyl-QA |
|  |  | CM30 | 501.1402_13.83 | C<sub>25</sub>H<sub>25</sub>O<sub>11</sub> | Cinnamic acid, Hex |
|  |  | CM15 | 711.2289_10.48 | C<sub>36</sub>H<sub>39</sub>O<sub>15</sub> | FA-Hex-Hex |
|  |  | CM17 | 401.1084_8.67 | C<sub>17</sub>H<sub>21</sub>O<sub>11</sub> | QA |
|  |  | CM28 | 373.1135_4.98 | C<sub>16</sub>H<sub>21</sub>O<sub>10</sub> | Hex |
|  |  | CM34 | 711.2289_8.67 | C<sub>36</sub>H<sub>39</sub>O<sub>15</sub> | Hex-FA-Hex |
|  |  | CM13 | 707.1976_5.71 | C<sub>36</sub>H<sub>35</sub>O<sub>15</sub> | CA, Hydroxy-FA-CA |
|  |  | CM32 | 529.1351_12.28 | C<sub>26</sub>H<sub>25</sub>O<sub>12</sub> | 3-CA, 4-FA or 4-FA, 5-CA |
|  |  | CM33 | 529.1351_11.94 | C<sub>26</sub>H<sub>25</sub>O<sub>12</sub> | 3-FA, 4-CA or 3-FA, 5-CA |
| Flavonoids |  | CM1 | 283.0606_14.01 | C<sub>16</sub>H<sub>11</sub>O<sub>5</sub> | / |
|  |  | CM2 | 283.0606_15.24 | C<sub>16</sub>H<sub>11</sub>O<sub>5</sub> | / |
|  |  | CM3 | 283.0606_19.92 | C<sub>16</sub>H<sub>11</sub>O<sub>5</sub> | / |
|  |  | CM9 | Linarin | C<sub>28</sub>H<sub>31</sub>O<sub>14</sub> | 7-*O*-Hex-dHex |
|  |  | CM4 | 313.0712_12.91 | C<sub>17</sub>H<sub>13</sub>O<sub>6</sub> | CH<sub>3</sub>, CH<sub>3</sub> |
|  |  | CM11 | 609.1244_12.06 | C<sub>30</sub>H<sub>25</sub>O<sub>14</sub> | CA-Hex |
|  |  | CM20 | 533.1448_17.17 | C<sub>29</sub>H<sub>25</sub>O<sub>10</sub> | Rupestonic acid |
|  |  | CM27 | 313.0712_12.02 | C<sub>17</sub>H<sub>13</sub>O<sub>6</sub> | CH<sub>3</sub>, CH<sub>3</sub> |
|  |  | CM5 | 313.0712_18.25 | C<sub>17</sub>H<sub>13</sub>O<sub>6</sub> | CH<sub>3</sub> |
|  |  | CM10 | 607.1663_12.03 | C<sub>28</sub>H<sub>31</sub>O<sub>15</sub> | CH<sub>3</sub>, 3-*O*-Pen-Hex |
|  |  | CM19 | 489.1186_12.97 | C<sub>27</sub>H<sub>21</sub>O<sub>9</sub> | CH<sub>3</sub>, 3-*O*-GlcUA |
|  |  | CM24 | 651.1567_11 | C<sub>29</sub>H<sub>31</sub>O<sub>17</sub> | CH<sub>3</sub>, 3-*O*-Hex-GlcUA |
|  |  | CM25 | 653.1659_12 | C<sub>36</sub>H<sub>29</sub>O<sub>12</sub> | CH<sub>3</sub>, HCA-FA |
|  |  | CM26 | 769.1827_8.54 | C<sub>33</sub>H<sub>37</sub>O<sub>21</sub> | 3-*O*-Hex-Pen-GlcUA |
|  |  | CM12 | 653.1506_9.71 | C<sub>32</sub>H<sub>29</sub>O<sub>15</sub> | CH<sub>3</sub>, CH<sub>3</sub>, CA-Hex |
|  |  | CM21 | 547.1246_12.59 | C<sub>29</sub>H<sub>23</sub>O<sub>11</sub> | CH<sub>3</sub>, CH<sub>3</sub>, Propylcaffeate |
|  |  | CM14 | 711.1773_8.66 | C<sub>31</sub>H<sub>35</sub>O<sub>19</sub> | Butanoic acid, 3-*O*-Hex-Hex |
|  |  | CM22 | Rutin | C<sub>27</sub>H<sub>30</sub>O<sub>16</sub> | 3-*O*-Hex-dHex |
|  |  | CM18 | 477.1038_7.2 | C<sub>22</sub>H<sub>21</sub>O<sub>12</sub> | CH<sub>3</sub>, 3-*O*-Hex |
|  |  | CM31 | 519.1291_13.17 | C<sub>28</sub>H<sub>23</sub>O<sub>10</sub> | CH<sub>3</sub>, 3-*O*-Ethane-FA |
|  |  | CM8 | 577.1557_13.02 | C<sub>27</sub>H<sub>29</sub>O<sub>14</sub> | 3-*O*-Pen-Hex |
|  |  | CM23 | 623.1612_11.81 | C<sub>28</sub>H<sub>31</sub>O<sub>16</sub> | 3-*O*-Hex-GlcUA |
|  |  | CM7 | 419.0978_13.15 | C<sub>20</sub>H<sub>19</sub>O<sub>10</sub> | Methoxy-phenylacetic acid |
|  |  | CM16 | 393.0827_7.61 | C<sub>22</sub>H<sub>17</sub>O<sub>7</sub> | OH, OCH<sub>3</sub>, *O*-ethylene glycol |

**Table 1.** Structural identification of 34 CMs in artificial and wild *Artemisia rupestris* L.
|  | Basic structure | Chemical markers | *m/z*_Rt | Elemental composition | Substituent groups |
| --- | --- | --- | --- | --- | --- |
| Chlorogenic acid/Caffeic acid derivatives |  | CM6 | 355.1029_6.54 | C<sub>16</sub>H<sub>19</sub>O<sub>9</sub> | Hex |
|  |  | **CM29** | 423.0927_6.81 | C<sub>19</sub>H<sub>19</sub>O<sub>11</sub> | Acetic acid alkenyl-QA |
|  |  | **CM30** | 501.1402_13.83 | C<sub>25</sub>H<sub>25</sub>O<sub>11</sub> | Cinnamic acid, Hex |
|  |  | **CM15** | 711.2289_10.48 | C<sub>36</sub>H<sub>39</sub>O<sub>15</sub> | FA-Hex-Hex |
|  |  | **CM17** | 401.1084_8.67 | C<sub>17</sub>H<sub>21</sub>O<sub>11</sub> | QA |
|  |  | CM28 | 373.1135_4.98 | C<sub>16</sub>H<sub>21</sub>O<sub>10</sub> | Hex |
|  |  | **CM34** | 711.2289_8.67 | C<sub>36</sub>H<sub>39</sub>O<sub>15</sub> | Hex-FA-Hex |
|  |  | **CM13** | 707.1976_5.71 | C<sub>36</sub>H<sub>35</sub>O<sub>15</sub> | CA, Hydroxy-FA-CA |
|  |  | CM32 | 529.1351_12.28 | C<sub>26</sub>H<sub>25</sub>O<sub>12</sub> | 3-CA, 4-FA or 4-FA, 5-CA |
|  |  | CM33 | 529.1351_11.94 | C<sub>26</sub>H<sub>25</sub>O<sub>12</sub> | 3-FA, 4-CA or 3-FA, 5-CA |
| Flavonoids |  | CM1 | 283.0606_14.01 | C<sub>16</sub>H<sub>11</sub>O<sub>5</sub> | / |
|  |  | CM2 | 283.0606_15.24 | C<sub>16</sub>H<sub>11</sub>O<sub>5</sub> | / |
|  |  | CM3 | 283.0606_19.92 | C<sub>16</sub>H<sub>11</sub>O<sub>5</sub> | / |
|  |  | CM9 | Linarin | C<sub>28</sub>H<sub>31</sub>O<sub>14</sub> | 7-*O*-Hex-dHex |
|  |  | CM4 | 313.0712_12.91 | C<sub>17</sub>H<sub>13</sub>O<sub>6</sub> | CH<sub>3</sub>, CH<sub>3</sub> |
|  |  | CM11 | 609.1244_12.06 | C<sub>30</sub>H<sub>25</sub>O<sub>14</sub> | CA-Hex |
|  |  | **CM20** | 533.1448_17.17 | C<sub>29</sub>H<sub>25</sub>O<sub>10</sub> | Rupestonic acid |
|  |  | CM27 | 313.0712_12.02 | C<sub>17</sub>H<sub>13</sub>O<sub>6</sub> | CH<sub>3</sub>, CH<sub>3</sub> |
|  |  | CM5 | 313.0712_18.25 | C<sub>17</sub>H<sub>13</sub>O<sub>6</sub> | CH<sub>3</sub> |
|  |  | **CM10** | 607.1663_12.03 | C<sub>28</sub>H<sub>31</sub>O<sub>15</sub> | CH<sub>3</sub>, 3-*O*-Pen-Hex |
|  |  | **CM19** | 489.1186_12.97 | C<sub>27</sub>H<sub>21</sub>O<sub>9</sub> | CH<sub>3</sub>, 3-*O*-GlcUA |
|  |  | CM24 | 651.1567_11 | C<sub>29</sub>H<sub>31</sub>O<sub>17</sub> | CH<sub>3</sub>, 3-*O*-Hex-GlcUA |
|  |  | **CM25** | 653.1659_12 | C<sub>36</sub>H<sub>29</sub>O<sub>12</sub> | CH<sub>3</sub>, HCA-FA |
|  |  | **CM26** | 769.1827_8.54 | C<sub>33</sub>H<sub>37</sub>O<sub>21</sub> | 3-*O*-Hex-Pen-GlcUA |
|  |  | **CM12** | 653.1506_9.71 | C<sub>32</sub>H<sub>29</sub>O<sub>15</sub> | CH<sub>3</sub>, CH<sub>3</sub>, CA-Hex |
|  |  | **CM21** | 547.1246_12.59 | C<sub>29</sub>H<sub>23</sub>O<sub>11</sub> | CH<sub>3</sub>, CH<sub>3</sub>, Propylcaffeate |
|  |  | **CM14** | 711.1773_8.66 | C<sub>31</sub>H<sub>35</sub>O<sub>19</sub> | Butanoic acid, 3-*O*-Hex-Hex |
|  |  | CM22 | Rutin | C<sub>27</sub>H<sub>30</sub>O<sub>16</sub> | 3-*O*-Hex-dHex |
|  |  | CM18 | 477.1038_7.2 | C<sub>22</sub>H<sub>21</sub>O<sub>12</sub> | CH<sub>3</sub>, 3-*O*-Hex |
|  |  | **CM31** | 519.1291_13.17 | C<sub>28</sub>H<sub>23</sub>O<sub>10</sub> | CH<sub>3</sub>, 3-*O*-Ethane-FA |
|  |  | **CM8** | 577.1557_13.02 | C<sub>27</sub>H<sub>29</sub>O<sub>14</sub> | 3-*O*-Pen-Hex |
|  |  | **CM23** | 623.1612_11.81 | C<sub>28</sub>H<sub>31</sub>O<sub>16</sub> | 3-*O*-Hex-GlcUA |
|  |  | **CM7** | 419.0978_13.15 | C<sub>20</sub>H<sub>19</sub>O<sub>10</sub> | Methoxy-phenylacetic acid |
|  |  | **CM16** | 393.0827_7.61 | C<sub>22</sub>H<sub>17</sub>O<sub>7</sub> | OH, OCH<sub>3</sub>, *O*-ethylene glycol |

CA, caffeic acid; FA, ferulic acid; GlcUA, glycuronic acid; Hex, hexose; dHex, deoxyhexose;
Pen, pentose; QA, quinic acid; Rt, retention time

**Figure 5.**
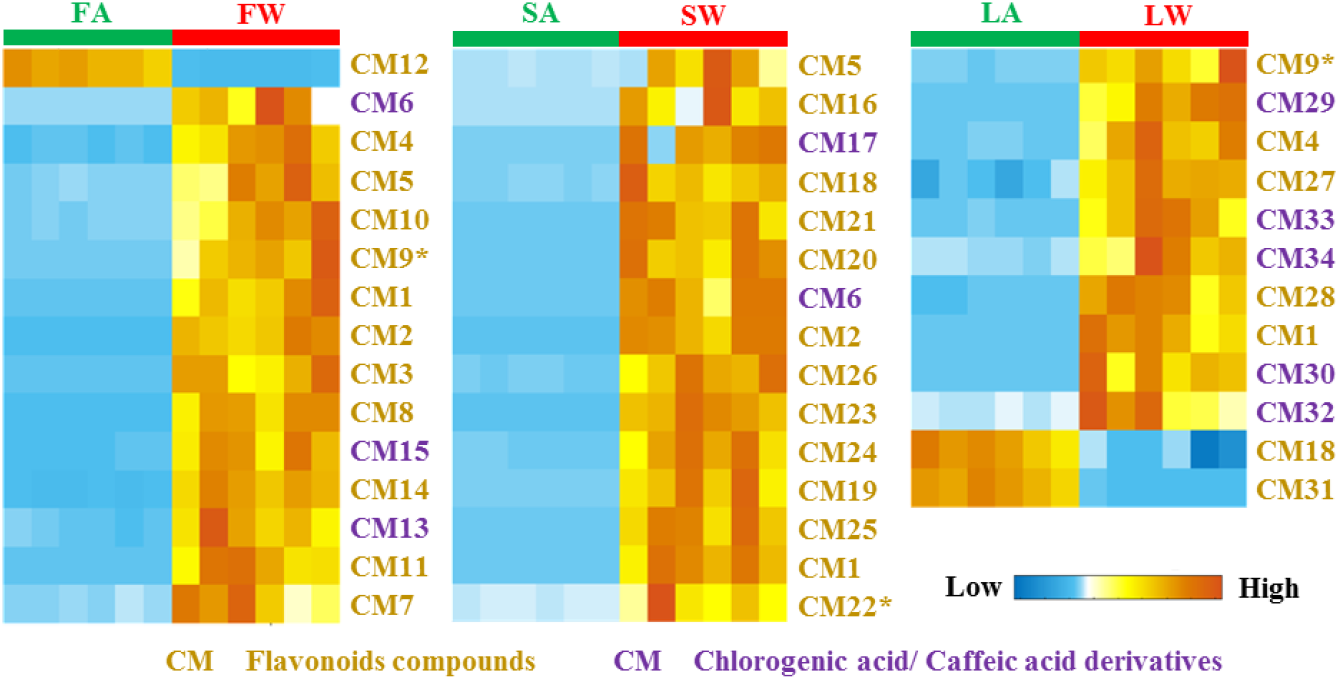
Cluster heatmap of CMs in artificial and wild *Artemisia rupestris* L. flower, stem, and leaf samples FA: flower, artificial group; FW: flower, wild group; SA: stem, artificial group; SW: stem, wild group; LA: leaf, artificial group; LW: leaf, wild group.

CM6, CM15, CM17, CM28, CM29, CM30, and CM34 showed prominent product ions at *m/z* 193, 149, and 134 in their MS/MS spectra, indicating a ferulic acid (FA) moiety in their structures. In particular, loss of 308.0901 Da in the CM30 MS/MS spectra was thought to occur via loss of a coumaroylhexoside moiety. CM15 exhibited a [M−H]^−^ ion at *m/z* 711, and weak ions at *m/z* 549 and *m/z* 387 arising from the sequential loss of two molecules. Furthermore, 162 Da was observed in the MS/MS spectrum, indicating that the two hexoses joined together. CM34 was an isomer of CM15; however, the ion at *m/z* 387 was absent in its MS/MS spectrum, thus the order of FA and hexose moieties in this metabolite was different from CM15.

CM13 displayed a [M−H]^−^ ion at *m/z* 707, and its MS/MS spectrum revealed a [quinic acid −H]^−^ ion at *m/z* 191 as the base peak, indicating that a quinic acid was substituted at the 3-OH or 5-OH position. Another weak ion at *m/z* 353 was inferred as caffeoylquinic acid according to the findings of a previous study (Gu et al. 2012). Thus, CM13 was identified as a quinic acid that was substituted by caffeoyl and hydroxyl-feruloyl-caffeoyl. A comparison of the MS/MS spectra of the two isomers, CM32 and CM33, showed that CM32 had base peak at *m/z* 173, which was inferred as 3-caffeoyl-4-feruoylquinic acid or 4-feruoyl-5-caffeoylquinic acid, while CM33 exhibited the highest ion at *m/z* 193, which was identified as 3-feruoyl-4-caffeoylquinic acid or 3-feruoyl-5-caffeoylquinic acid.

According to MS fragmentation regularity and characteristic product ions of different basic structures, eight flavones and 14 flavonol were putatively identified. For example, the characteristic product ions at *m/z* 283, 268, 240, 239, and 151 led to the aglycone identification as acacetin, including CM1, CM2, CM3,and CM9 (linarin). Metabolites with basic trihydroxydimethoxyflavone structures generated product ions at *m/z* 313, 298, and 283, such as CM5, CM10, and CM19. In addition, characteristic neutral losses also help structure analysis. Loss of 294 Da could be attributed to hexose (162 Da) and pentose (132 Da) groups, and 484 Da indicated the sugar unit may consist of a hexose (132 Da), a deoxyhexose (146 Da), and a glycuronic acid (176 Da).

The MS/MS spectrum of CM16 was distinctive in that had an ion *m/z* 299 as the highest peak and *m/z*151; however, there were no obvious characteristic product ions, such as *m/z* 284, 255 of methylkaempferol or other normal flavonoids. Gu et al. previously analyzed 2-phenoxychromones in *Artemisia rupestris* L., which display an [M−H]^−^ion at *m/z* 299. Thus, CM16 was putatively identified as hydroxyl-dihydroethoxy-2-phenoxychromone. The base peak in the MS/MS spectrum of CM7 was an ion at *m/z* 153, which indicated it was a type of chalcone.

### DEGs in flavonoids and chlorogenic acid/caffeic acid biosynthetic pathways

Flavonoids and CCDs belong to active substances of *Artemisia rupestris* L. Flavonoid biosynthesis derives from phenylpropanol metabolism (Supplementary Fig. S1), in which phenylalanine is converted to cinnamic acid by phenylalanine ammonia lyase (PAL). Cinnamic acid can then be used to generate caffeic acid, chlorogenic acid, and the skeleton structure of flavonoids. The present study focused on DEGs that had same function and variation trend in these pathways. In the W group flower (Fig. 6A), *PAL* gene expression was downregulated, indicating a reduced initial source of flavonoids and phenylpropanoids. This may have resulted in the downregulation of subsequent reactions, and our results showed that genes related to biosynthesis of caffeic acid, isoflavonoids, flavanone, and flavanonol, such as *DFR* and *CYP73A*, were downregulated. On the other hand, *CHS* and *F5H* were upregulated. Interestingly, the varietal discrepancy in the leaf samples was very different from that of the flower samples (Fig. 6B). It could be that the higher expression of *PAL* in wild *Artemisia rupestris* L. leaf led to an increased expression of many other key genes in subsequent pathways, including flavonoid hydroxylation (*F3H, CYP75A, CYP75B1*), flavonol synthesis (*FLS*), propionyl transference of isoflavone glycosides (*IF7MAT*), isoflavone methylation (*7-IOMT*), and chlorogenic acid synthesis (*C3’H*). Meanwhile, some genes involved in phenylpropanoid biosynthesis pathways were expressed at low levels, including *CAD* and *CCR*. The expression of DEGs in stem samples was in between that of flower and leaf samples. Elevated expression of *CHS* and *F5H* was consistent in the flower samples, and lower expression of *IF7GT* was similar in the leaf samples.

**Figure 6.**
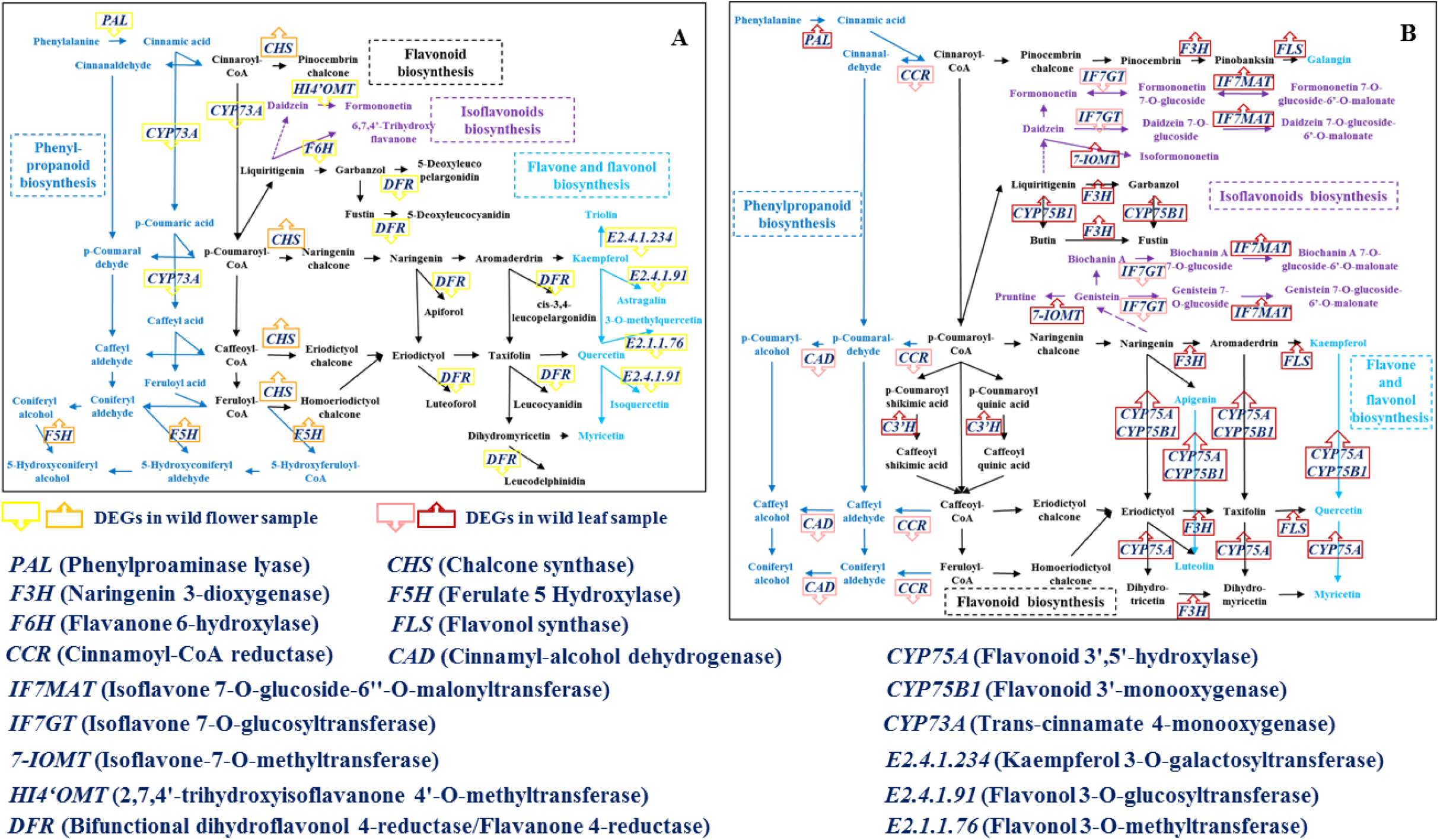
DEGs with explicit changes in direction in flavonoid and CCD biosynthetic pathways in *Artemisia rupestris* L. flower and leaf samples based on pathway enrichment analysis of DEGs.

### Genetic basis of CMs based on integration analysis and qRT-PCR

Correlation analysis between CMs and DEGs revealed that 13 DEGs showed a close relationship with 32 CMs in the three organ types (Supplementary Fig. S2). Subsequently, qRT-PCR was conducted to validate the reliability of these candidate DEGs. In wild flower samples (Fig. 7A), upregulation of Unigene98154_All, which could be *CHS*, showed a positive relationship with CM4 and CM7, downregulation of *DFR* showed a negative relationship with CM13, and *EC 2*.*4*.*1*.*92* or *EC 2*.*4*.*1*.*234* showed a positive relationship with CM12 and a negative relationship with CM5 and CM14. Although the other CMs were affected by multiple DEGs, their change in trend could be explained reasonably. In wild stem samples (Fig. 7B), it was confirmed that there was a much higher content of *F5H* than in the artificial stem, which was consistent with the increase ofCM19, CM24, and CM26. The increase in CM2, CM20, CM21, and CM25 could be explained by their negative response to downregulation of *IF7GT*. Under the dual function of *F5H* and *IF7GT*, the stronger effect from *F5H* resulted in the elevation of CM23 in the wild group. Finally, after validation, upregulation of *IF7MAT* was associated with increased (CM4, CM9, CM18, CM29, CM30, CM32 and CM34) and decreased (CM18) levels of eight CMs (Fig. 7C).

**Figure 7.**
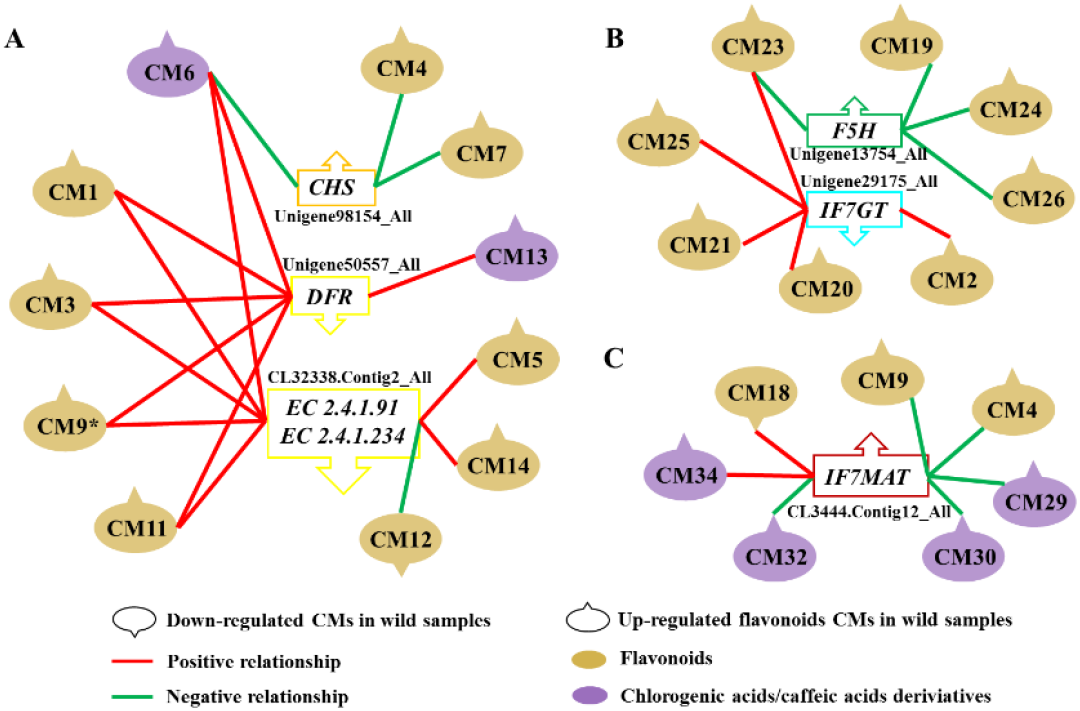
Confirmed relationships between CMs and DEGs by qRT-PCR.

## DISCUSSION

Plants grow in distinct environments may undergo remarkable reprogramming of their genes and metabolites. Thus, unraveling the effects of ecological factors at the molecular level is important for botanical research, especially for medicinal plants. Integration of metabolomics and transcriptomics is already used for comparative analyses of plants under different conditions (Wu et al., 2016; Sebastián et al., 2016; Xu et al., 2019; Wei et al., 2018; Guo et al., 2020). However, most of these have focused on one or several classes of known metabolites and model species and common crops. *Artemisia rupestris* L. is a phytomedicine whose material basis and active ingredients have not been fully studied. The untargeted and parallel strategy proposed in the present study using an integrated metabolomics–transcriptomics approach represents a useful high-throughput tool.

This research starting with untargeted metabolomics would help us obtain numerous differential species including primary and secondary metabolites. Besides, targeted attention to key known metabolites is recommended and enables the simplest understanding of the differences, as well as in case of important information omission during untargeted analysis. Emphatically, external validation was necessary for reliable markers, and could also reduce the workload of structure identification. *De novo* transcriptomics matched the tissue specificity of the metabolome and reference genome absence of *Artemisia rupestris* L. plants. Finally, the method and scale of the integration analysis could be selected focusing on the purpose of the study. The present study took advantage of the simplicity and intuitiveness of Pearson correlation analysis and, following qRT-PCR validation, discovered several novel and underexplored associations.

Our metabolomics results revealed clear differences between artificial and wild *Artemisia rupestris* L. plants. Among more than 300 differential varieties of each sample type, the FC values of at least 50 varieties were >10, the majority of which were flavonoids and CCDs. Importantly, with the exception of a few, many CMs showed much higher content in wild samples regardless of the organ. However, the expression of DEGs in the three organs was very different as most of the upregulated DEGs associated with CMs were present in the leaf samples. This indicated that the leaf may the dominant synthetic site of flavonoids and CCDs, and high concentrations of these metabolites could still accumulate in flowers and stems, although some biosynthetic pathways were repressed.

The conjugated structure of flavonoid compounds could protect the organism from harm of ultraviolet (UV) radiation, and a close relationship has been reported between flavonoids and light. Accumulation of flavonoids could be triggered or amplified by UV (Neugart et al. 2019), and compared with visible light, shorter wavelength light treatment would result in a significantly higher total flavonoid content (Liu et al. 2018). Chlorogenic acid plays an important role in the physiological resistance of plants. For example, the phenolic hydroxyl group present in their structure protects the organism from reactive oxygen species and free radicals, and could reduce UV damage and increase resistance to microorganisms (Tegelberg, et al. 2004; Clé et al. 2008). In the present study, wild *Artemisia rupestris* L. plants grew at a much higher altitude than the artificial variety. Thus, exposure of the wild variety to stronger UV radiation could have led to a higher content of flavonoids and CCDs. Moreover, it was reported that UV light affects plant morphology, and decreases the plant height and leaf area (Yan et al. 2019). The wild plants we collected were smaller than that of the artificial variety, and they also had short, negligible branches, and much denser leaves (Supplementary Fig. S3). The characteristics of the wild species contributed to facilitating flavonoid and CCD synthesis and efficient transport of these compounds to other parts of the plant to adapt to living in a wild environment. As expected, unigenes in “plant-pathogen interaction” and “circadian rhythm” belonging to environmental adaptation pathways were also upregulated in wild species.

The characteristic metabolite rupestonic acid in artificial *Artemisia rupestris* L. plant showed no obvious difference from that in the wild variety, and was slightly higher than the wild variety. The other rupestonic acid derivatives were the similar. According to the findings of our previous study, rupestonic acid and its derivatives were mainly distributed in the flower. Our transcriptome results confirmed that the key DEGs involved in the terpenoid backbone biosynthesis pathway were downregulated in the wild flower samples. These data indicate that the ecological environment had little influence on rupestonic acid and its derivatives, and artificial varieties could be used for drug development directing at this category of ingredients.

In summary, our study was the first time to compare artificial and wild *Artemisia rupestris* L. from the level of metabolome and transcriptome in a systematic manner, and 34 reliable chemical markers as well as their genetic basis was discovered. During this process, 19 potential new metabolites were identified, which illustrated high throughput of the strategy used in this study. The results could provide novel molecular information for material basis research and quality control of *Artemisia rupestris* L., and also provide reference of research methods and ideas for other phytomedicines.

## MATERIALS AND METHODS

### Plant materials

Artificially cultivated and wild varieties of *Artemisia rupestris* L. were collected at the full-bloom stage in the Altay–Fuyun region, Xinjiang, China at the altitude of about 800 meters. Fresh samples were immediately frozen in liquid nitrogen and then transferred to −80 °C for long-term storage. Flower, stem, and leaf samples of *Artemisia rupestris* L. were analyzed separately.

### Chemicals and reagents

Chrysosplenetin B was purchased from Chroma Biotechnology Company Limited (China). Linarin, vanillic acid, luteolin-7-*O*-glucuronide, and chlorogenic acid were purchased from the National Institutes for Food and Drug Control (China). Vitexicarpin, rutinum, rupestonic acid, luteolin, isoquercitin, and artemitin were purchased from Chenguang Biotechnology Company Limited (China). Taxifolin,6-hydroxyquercetin-7-*O*-β-_D_-glucopyranoside, kaempferol, pinocembrin, apigenin, isohamnetin, caffeic acid, syringic acid, cacetin, kaempferide, engeletin, baicalin, and hyperoside were purchased from ConBon Biotech Company Limited (China). Lysionotin was purchased from SenBeijia Biological Technology Company Limited (China). The purities of the above 25 compounds were >98%.

Methanol and acetonitrile (HPLC-grade) were purchased from Merck (Germany). Formic acid (HPLC-grade) was obtained from Roe Scientific Inc. (USA). DEPC water was purchased from Ambion (USA). Chloroform, β-mercaptoethanol, isoamyl alcohol, and isopropyl alcohol were purchased from Xilong Chemical Co. Ltd (China).

### Metabolome extraction

Freeze-dried samples were ground to a powder and each 50-mg aliquot was extracted by adding of 1 mL methanol-water solvent (8:2, *v/v*, stored at 4 °C) followed by high-speed dispersion for 3 min (25,000 rpm for 1 min, 3,000 rpm for 1 min, 25,000 rpm for 1 min). After ultrasound treatment for 10 min, the mixture was centrifuged at 13,500 *rpm* at 4 °C for 10 min. The supernatant was collected and evaporated to dryness in a SpeedVac concentrator (Thermo Savant). Sample residues were reconstituted in 1 mL acetonitrile-water (1:1, *v/v*) and mixed for 10 min by ultrasonic treatment. After centrifugation at 13,500 rpm at 4 °C for 1 min, the supernatants were filtered using a 0.22-µm microporous membrane for LC–MS analysis.

### LC–MS conditions

Chromatographic separation was performed on a reversed-phase Waters HSS T3 column (100 mm × 2.1 mm, 1.8 μm) using an ultra-performance LC system (UltiMate 3000; Thermo Fisher Scientific).The column was maintained at 40 °C and flushed with aqueous 0.1% formic acid (A) and acetonitrile 0.1% formic acid (B) using a flow rate of 300 μL/min. The gradient conditions were as follows: 95%–50% (v/v) B at 0–20 min, 50%–2% B at 20–27 min, 2%–2% B at 27–30 min. The injection volume was 5 μL.

MS was performed using a Q-OT-qIT hybrid mass spectrometer (Orbitrap Fusion Lumos, Thermo Fisher Scientific, USA) equipped with an ESI source. Data were acquired in negative ion mode. The instrumental parameters were as follows: spray voltage, −3.2 kV; vaporizer temperature, 350 °C; capillary temperature, 350 °C; resolution, 60,000; and automatic gain control target, 1.0e^6^. The scan range was *m/z* 100–1000. The PRM method in the validation assay was built according to the retention time, exact mass, and optimized collision energy of targeted metabolites. Data were acquired using Xcalibur 4.0 software.

Samples were injected in a random order. QC samples derived from different organs were prepared by mixing equal aliquots from all specific types of authentic samples, termed flower-QC, stem-QC, and leaf-QC. QC samples were inserted and analyzed regularly per 12 authentic samples to monitor the stability of the LC–MS system (Gika et al., 2007).

### Data analysis of metabolomics

Raw data were converted to mzXML format using the MassMatrix MS Data File Conversion Tools (http://www.massmatrix.net). Peak detection and alignment were then performed using XCMS package of R software (version 3.15.2; R project, Vienna, Austria). SIMCA-P (version 15.0) software was used for multivariate statistical analysis of the data obtained via UHPLC–MS analysis. A PCA model was used to obtain an overview of the distribution of samples and identify outliers. The OPLS-DA model was used to identify differential metabolites contributing to group clustering. Finally, potential biomarkers were identified according to their exact *m/z* value, MS/MS spectrum, and retention times. METLIN (http://metlin.scripps.edu/) was used for database searching.

### RNA isolation and sequencing

Total RNA from each 30-mg aliquot of flower, stem, and leaf samples were isolated using 1.5 mL 2% CTAB lysis solution including 2% β-mercaptoethanol. After incubation at 1000 rpm at 65 °C for 20 min, the supernatants were treated twice with chloroform-isoamyl alcohol solvent (24:1, *v/v*) and isopropyl alcohol. The sediment was then washed using 75% ethyl alcohol and the dried RNA was dissolved using DEPC water for subsequent analysis. The quality and quantity of RNA were measured using an Agilent 2100 Bioanalyzer (Agilent Technologies, USA) and Nanodrop spectrophotometer (NanoDrop 8000, Thermo Fisher Scientific, USA), respectively. RNA samples with an RNA integrity number >7 were sent to the Rockefeller University Genomics Resource Center (New York, USA) for next-generation sequencing using IlluminaHiSeq.

### De novo transcriptome assembly

Raw data obtained after sequencing were filtered reads that contained adapters, >10%unknown nucleosides, and >50% low quality bases (Q<20). Clean reads were then assembled into unique consensus sequences using Trinity. The redundancy was eliminated using Tgicl and further assembled into a single set of non-redundant unigenes.

### Functional annotation and differential pathway analysis

Unigenes were annotated using BLASTX (E-value cutoff of 10^−5^) against six databases: National Centre for Biotechnology Information (NCBI) non-redundant (Nr), NCBI nucleotide (nt), Gene Ontology (GO), Cluster of Orthologous Groups (COG), Swiss-Prot, and Gene ontology and Kyoto Encyclopedia of Genes and Genomes (KEGG).Using the transcriptome assembled by Trinity as a reference, clean reads of each sample were mapped using RSEM. The normalized read counts from three replicates of each sample were analyzed, and unigenes that had significant differences in expression were determined using DEseq. The FDR *q* value threshold was set to 0.005, and the fold change in expression was set to 2.0. KEGG enrichment analysis was then performed using KOBAS 2.0 with hypergeometric tests to determine the distribution of unigenes based on their biological pathways. Pathways with FDR *q* values ≤0.05 were considered to be significantly enriched.

### qRT-PCR

The expression of selected DEGs was determined by qRT-PCR. Gene-specific primers were designed for 13 candidate genes based on obtained data. PCR was performed at 56 °C and 94 °C for 3 min 45 s, respectively. The thermal cycling conditions were as follows: 40 cycles at 94 °C for 20 s for denaturation, and 56 °C for 20 s and 72 °C for 20 s for annealing and extension. After these reactions, dissociation curve analysis was performed to evaluate the specificity of the primers. The housekeeping gene, β-tubulin, which has highly abundant and stable expression, was used as reference gene. All reactions in all experiments were repeated three times.

## SUPPLEMENTAL DATA

The following supplemental materials are available.

**Supplemental Table S1**. Detail information of known representative metabolites in Artemisia rupestris L.

**Supplemental Table S2**. Assembly quality statistics of the Artemisia rupestris L. transcriptome.

**Supplemental Table S3**. MS/MS data of chemical markers that discovered in three organ types of artificial and wild *Artemisia rupestris* L.plants.

**Supplemental Figure S1**. Diagram of synthesis of flavonoid and chlorogenic acids. **Supplemental Figure S2**. Morphological feature comparison of artificial and wild Artemisia rupestris L. a) artificial variety; b) wild variety.

## ACKNOWLEDGEMENT

This work was funded by the Natural Science Foundation of China (Grant No. 2167050718 and Grant No. 82003714). We also thank Liu Geyu and Aisa Haji Akber working in the Key Laboratory of Xinjiang Indigenous Medicinal Plants Resource Utilization, Xinjiang Technical Institute of Physics and Chemistry, Chinese Academy of Sciences for their support in sample collection.

